# Deep learning for brains?: Different linear and nonlinear scaling in UK Biobank brain images vs. machine-learning datasets

**DOI:** 10.1101/757054

**Authors:** Marc-Andre Schulz, B.T. Thomas Yeo, Joshua T. Vogelstein, Janaina Mourao-Miranada, Jakob N. Kather, Konrad Kording, Blake Richards, Danilo Bzdok

**Affiliations:** Department of Psychiatry, Psychotherapy, and Psychosomatics, Rheinisch-Westfälische Technische Hochschule (RWTH) Aachen University, Aachen, Germany; Department of Electrical and Computer Engineering, ASTAR-NUS Clinical Imaging Research Centre, Singapore; Institute for Neurotechnology and Memory Networks Program, National University of Singapore, Singapore; Centre for Cognitive Neuroscience, Duke-NUS Medical School, Singapore; Department of Biomedical Engineering, Institute for Computational Medicine, Johns Hopkins University, Baltimore, USA; Kavli Neuroscience Discovery Institute, Johns Hopkins University, Baltimore, USA; Max Planck University College London Centre for Computational Psychiatry and Ageing Research, University College London, UK; Centre for Medical Image Computing, Department of Computer Science, University College London, UK; Department of Medicine III, University Hospital RWTH Aachen, Aachen, Germany; German Cancer Consortium (DKTK), German Cancer Research Center (DKFZ), Heidelberg, Germany; Applied Tumor Immunity, German Cancer Research Center (DKFZ), Heidelberg, Germany; Department of Neuroscience and Department of Bioengineering, University of Pennsylvania, Philadelphia, Pennsylvania; Department of Neurology and Neurosurgery, McGill University, Montréal, Québec, Canada; School of Computer Science, McGill University, Montréal, Québec, Canada; Canadian Institute for Advanced Research, Toronto, Ontario, Canada; Mila, Montréal, Québec, Canada; Neurospin, Commissariat à l’Energie Atomique (CEA) Saclay, Gif-sur-Yvette, France; Parietal Team, Institut National de Recherche en Informatique et en Automatique (INRIA), France

## Abstract

In recent years, deep learning has unlocked unprecedented success in various domains, especially in image, text, and speech processing. These breakthroughs may hold promise for neuroscience and especially for brain-imaging investigators who start to analyze thousands of participants. However, deep learning is only beneficial if the data have nonlinear relationships and if they are exploitable at currently available sample sizes. We systematically profiled the performance of deep models, kernel models, and linear models as a function of sample size on UK Biobank brain images against established machine learning references. On MNIST and Zalando Fashion, prediction accuracy consistently improved when escalating from linear models to shallow-nonlinear models, and further improved when switching to deep-nonlinear models. The more observations were available for model training, the greater the performance gain we saw. In contrast, using structural or functional brain scans, simple linear models performed on par with more complex, highly parameterized models in age/sex prediction across increasing sample sizes. In fact, linear models kept improving as the sample size approached ∼10,000 participants. Our results indicate that the increase in performance of linear models with additional data does not saturate at the limit of current feasibility. Yet, nonlinearities of common brain scans remain largely inaccessible to both kernel and deep learning methods at any examined scale.

## Introduction

Following genetics and other biological domains, imaging neuroscience recently started to become a “big data” science (Efron 2012; Thompson et al. 2017; Smith & Nichols 2018). The brain sciences have been proposed to be one of the most data-rich medical specialties (Nature Editorial 2016) due to amassing high-resolution imaging data.

Several data collection initiatives stand out in the brain-imaging landscape (Smith & Nichols 2018), including the Human Connectome Project (HCP), the UK Biobank (UKBB) imaging Study, and the Enhancing NeuroImaging Genetics through Meta-Analysis (ENIGMA) Consortium. The UKBB is perhaps the most compelling, as this resource includes genetic profiling and an extensive variety of phenotyping descriptors. The data aggregation set out in 2006 to gather genetic and environmental data of 500,000 volunteers and is currently the world’s largest biomedical dataset (Sudlow et al., 2015). The participants will be followed for >25 years, including repeated measurements and full access to electronic health records. In 2014 UKBB launched its brain-imaging extension, aiming to gather several magnetic resonance imaging (MRI) modalities of 100,000 participants by 2022 (Miller et al. 2016). UKBB is specifically designed for prospective population epidemiology. Instead, the ambition of HCP lies in functional and anatomical connectivity in healthy subjects, whereas ENIGMA places an important emphasis on genetic profiling in combination with brain scanning in psychiatric and neurological disease. The creation, curation, and collaboration of large-scale brain-imaging datasets with thousands of subjects promises to enable more ambitious statistical analyses than are currently the norm (Bzdok & Yeo, 2017).

An important benefit of such large-scale data collections is that they may allow for more expressive models that could more powerfully isolate and describe phenomena in the brain - models that can capture complicated nonlinear interactions dormant in common, abundant brain scans. Spurred by increasing data availability, analysis of brain-imaging data is more and more often resorting to sophisticated machine learning algorithms (Bzdok 2017; Marquand et al. 2016; Marblestone et al. 2016). A core step in data analysis has always been the identification of the most relevant variables to be included and how they should be encoded or built into newly designed features, often called “manual feature engineering” (Abu-Mostafa et al. 2012).

Linear models have long dominated data analysis, as complex transformations into rich high-dimensional spaces were historically computationally infeasible. Towards the end of the 20th century, kernel embeddings (Schölkopf et al. 2002) were devised to efficiently map data to rich high-dimensional spaces in a computationally efficient fashion. Specifically, kernel methods rely on the pairwise similarities between data points instead of explicitly computing coordinates of all data points in high-dimensional space. Kernel methods can thus perform data analysis within an enriched, potentially infinitely dimensional representation of the original input variables. This extension of many classical linear methods towards capturing more complicated nonlinear patterns in data enabled enhanced prediction accuracy in a large variety of applications, including many areas of biomedicine (Borgwardt 2011).

“Preprogrammed” kernel methods operating on predefined similarity functions, in turn, have recently been superseded by the renaissance of artificial neural networks under the term “deep learning” (LeCun et al. 2015; Goodfellow et al. 2016; Schmidhuber 2015). One key aspect of this even more flexible class of methods is the cascade of successive nonlinear transformations from the input variables. In natural images, for instance, pixel inputs provide color information for each single location. A deep neural network automatically learns to combine these pixels into basic shapes, like circles or edges, which get combined by further transformation layers into objects like furniture, which eventually compose whole scenes and other abstract concepts, like a kitchen, represented in the highest layers. Going beyond kernel methods, deep methods have enabled “automatic feature engineering” and even richer representations and abstractions of patterns in data. In a sense, deep learning methods can be thought of as kernel methods that also learn the kernel (Giryes et al. 2015).

This upgrade has unlocked unprecedented prediction success in a number of application domains, especially those involving the processing of natural images, text, or speech - areas where deep neural networks can leverage their strengths: Compositional representations and a hardcoded assumption of translation invariance. Whether deep learning will be equally successful in brain-imaging, specifically in predicting phenotypes from structural and functional MRI, remains yet unclear (He et al. 2018) and a systematic evaluation of deep learning in brain imaging is urgently required. However, deep learning models are highly flexible and new varieties are constantly developed. This rank growth of deep learning models makes it near impossible to comprehensively benchmark deep learning for brain-imaging. Here, we address this need by first principles and investigated the precondition most likely to limit the success of deep learning on brain-imaging data: The extent to which nonlinear relationships in brain images are exploitable at currently available sample sizes.

In short, kernel-based models have outperformed linear methods in many applications. Deep-learning models have again refined pattern extraction and thus further boosted prediction accuracy in structured data such as natural images. This study investigated the scaling behaviors of these three modeling regimes on brain-imaging data and compared it to the performance profiles on standard machine learning datasets. We thus provide some first answers to the question: “Do emerging large-scale brain-imaging datasets contain nonlinear information that can be exploited by currently available kernel and deep models?”

## Results

Our goal was to assess how much the analysis of brain-imaging data can benefit from using nonlinear methods or even deep learning. For results to be generalizable, we wished to shed light on the general types of information in brain-imaging data, and whether those properties demand more complicated models making the most out of increasing sample size. Gaining such important intuitions would allow us to not only observe that a certain approach works well on brain-imaging data, but draw conclusions why that might be the case.

To this end, we evaluated how the prediction performance scales with increasing sample sizes for model classes with increasing prediction capacity and datasets of increasing prediction difficulty. We considered the achieved prediction performance as a function of the available sample size to obtain a principled empirical assessment of sample complexity (J. Friedman et al. 2001; Goodfellow et al. 2016; Abraham et al. 2017). Analyzing the empirical sample complexity allows for insight into the information content of data as perceived through the assumptions of a given model class. For example, linear models are blind to nonlinear patterns in data by construction. Nonlinear kernel methods assume that a certain type of nonlinear interaction (defined by the “kernel”) is best suited to identify decision boundaries for accurate classification. Going one step further, deep neural networks also expect nonlinear interactions in the data, but, intuitively, they learn the kernel, rather than assuming it.

We used three classes of learning algorithms to evaluate cross-validated prediction performance on reference datasets (Fig. 1): a) Classical linear models are used for a number of decades since the middle of the 20th century in various empirical domains (Tikhonov 1963; Tikhonov et al. 2013), b) kernel support vector machines that were among the most competitive approaches from the late 1990s to approximately 2010 and c) common deep neural network algorithms that have now come to dominate areas where powerful empirical predictions are key and large amounts of structured (e.g., images) training data is available. For each model class we chose three representative algorithms, attempting to cover the plurality of approaches in a given regime of quantitative analysis. For the class of linear methods, we chose linear discriminant analysis (LDA), logistic regression, and linear support vector machines (SVMs). For shallow-nonlinear models, we profiled extensions of linear models with the popular radial-basis, polynomial, and sigmoidal kernels. For deep-nonlinear models, we used fully-connected and convolutional deep neural networks with and without GAP. We thus juxtaposed several analysis methods that recapitulate important periods in the recent data science evolution.

**Fig. 1:**
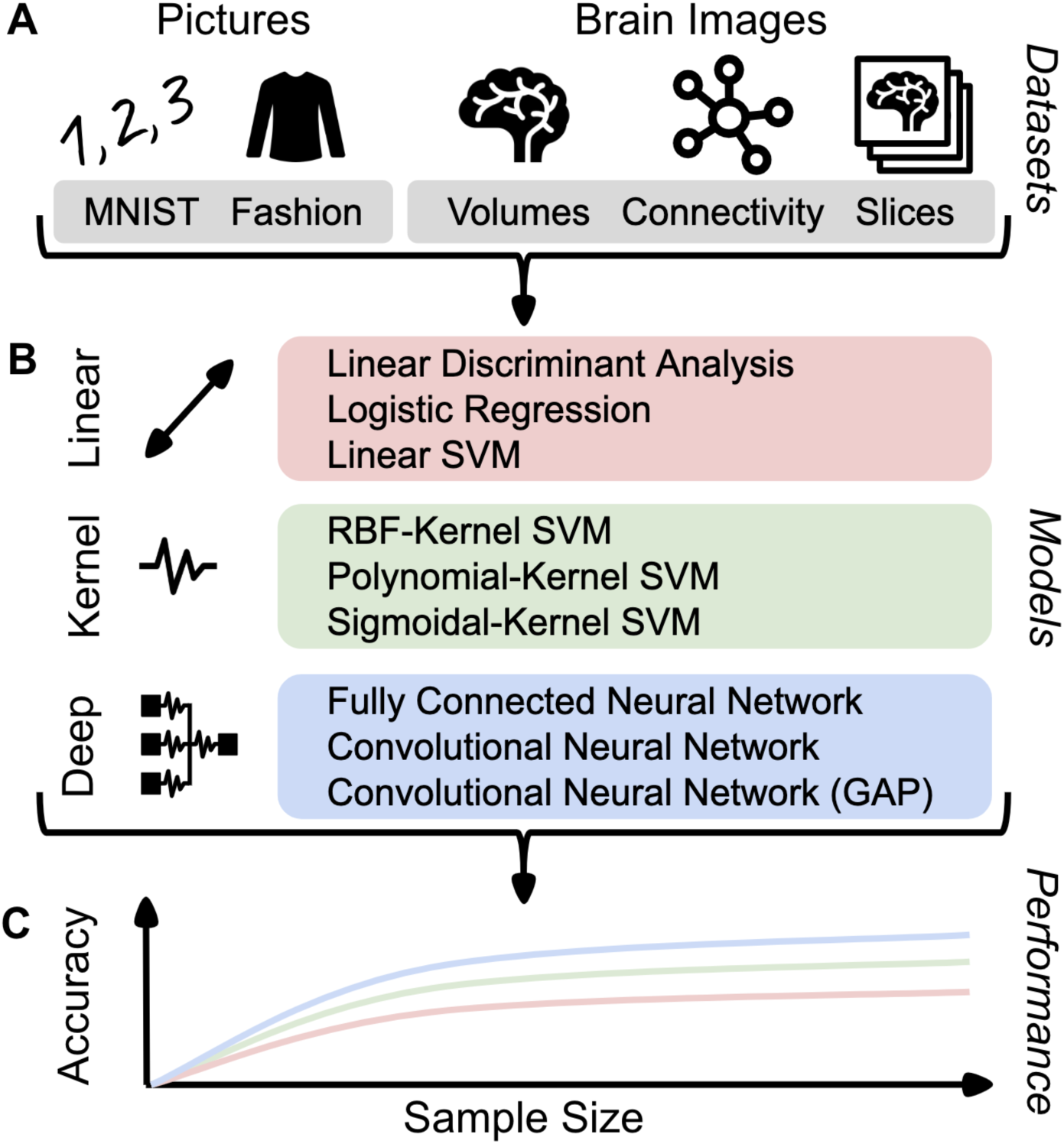
Workflow and experimental design. Properties of machine-learning datasets were directly compared to brain-imaging datasets based on performance curves of three classes of predictive models. A) Well-understood benchmark datasets from the machine learning community were selected to replicate previous results and serves as a reference point for the scaling behavior to imaging neuroscience: The MNIST images were to be classified into handwritten digits and Zalando’s Fashion images were to be classified into types of clothing. Several representations of brain structure and function were obtained from the UK Biobank resource: Region volumes and whole-brain slices from structural MRI, as well as functional connectivity strengths from resting state functional MRI. Brain image data were used to predict the subjects’ age and sex in 10 subgroups to match the 10 target categories of MNIST and Fashion. B) Prediction performance for each dataset was profiled using predictive models that have capacity for increasing predictive power: linear models (red tone), shallow-nonlinear kernel SVMs (green tone), and deep-nonlinear neural network algorithms (blue tones). C) For each combination of dataset and model, we systematically varied the number of data points available for the training the model. The resulting empirical estimates of sample complexity allow to extrapolate conclusions to always larger sample sizes.

### Model performance on machine learning reference datasets

To verify that we can obtain empirical estimates of the sample complexity of linear, kernel, and deep models, we initially examined two reference datasets that have been pervasively used in the machine learning community. This would not only validate our approach to chart the scaling behavior of predictive performance with increasing sample size, but also establish a point of comparison for performance differences between the three model classes and the brain-imaging data. The MNIST dataset (Modified National Institute of Standards and Technology dataset; LeCun & Cortes 2010) is an important classical benchmark datasets in the machine learning community. This resource consists of greyscale images of handwritten digits with the digit value (‘0’-’9’) to be classified from the raw pixel information. To quantitatively characterize the effects of a more challenging prediction goal, we also analyzed Zalando’s Fashion dataset (“Fashion-MNIST”, Xiao et al. 2017). The sometimes called “Fashion-MNIST” dataset consists of greyscale images of fashion products, for classification of ten categories of clothing. This resource was created to provide a more difficult prediction problem than MNIST while retaining the same number of targets (i.e., 10 categories), dimensionality (i.e., 28×28 = 768 variables), and sample size (i.e., 70,000 images).

To verify that our model implementations work as would be expected from the technical literature, we replicated several current-best prediction performances on the MNIST dataset. On the full dataset comprising 60,000 training observations, our logistic regression model achieved 91.32±0.90 % classification performance (out-of-sample prediction accuracy mean±standard deviation across resampling iterations), which is comparable to earlier work <u>(91.7% reported in Xiao et al. 2017)</u>. The more modern support vector machine with a radial-basis-function kernel (RBF-SVM) achieved 96.79±0.65 % accuracy (98.6% in LeCun & Cortes 2010). Finally, standard convolutional deep neural network model achieved 99.03±0.39 % prediction accuracy - in line with the 98.9% reported by LeCun et al. (1998) based on a similar deep learning architecture. Our model performances replicated state-of-the-art classification accuracies that were reported in previous research in the technical communities.

To carefully quantify the degree to which nonlinear structure can be exploited in MNIST and Fashion datasets, we compared the performance of linear models and nonlinear kernel support vector machines with gradually increasing number of training images. Systematic improvements in prediction accuracy when moving from linear models to nonlinear models consistently indicated the existence of exploitable nonlinear information predictive of the digit category in the dataset. On the MNIST dataset (Fig. 2-A.1), linear and kernel models performed indistinguishably for low sample sizes. Yet, we observed that linear models and kernel models have diverged in prediction accuracy starting at around 1,000 example images. Exceeding this sample size, the worst among the three kernel models (RBF-SVM) outperformed the best among the three linear models (logistic regression). In the Fashion dataset (Fig. 2-A.2), the worst kernel model (sigmoid-SVM) began to outperform the best linear model (logistic regression) starting from 4,000 observations. This difference in performance scaling grew to 4.03 percentage points (p.p.) and 1.76 p.p. at 8,000 observations for MNIST and Fashion, respectively. That is, we show a less prominent gap between linear and shallow nonlinear methods in one of two comparable datasets with a more complicated prediction goal (i.e., detect clothing rather than handwritten digits). For sufficiently large sample sizes available for model training, kernel methods outperformed linear models in both the MNIST and Fashion datasets. Thus, kernel support vector machines could readily exploit nonlinear structure in the examined datasets that is inaccessible to simpler linear models by principle.

**Fig. 2:**
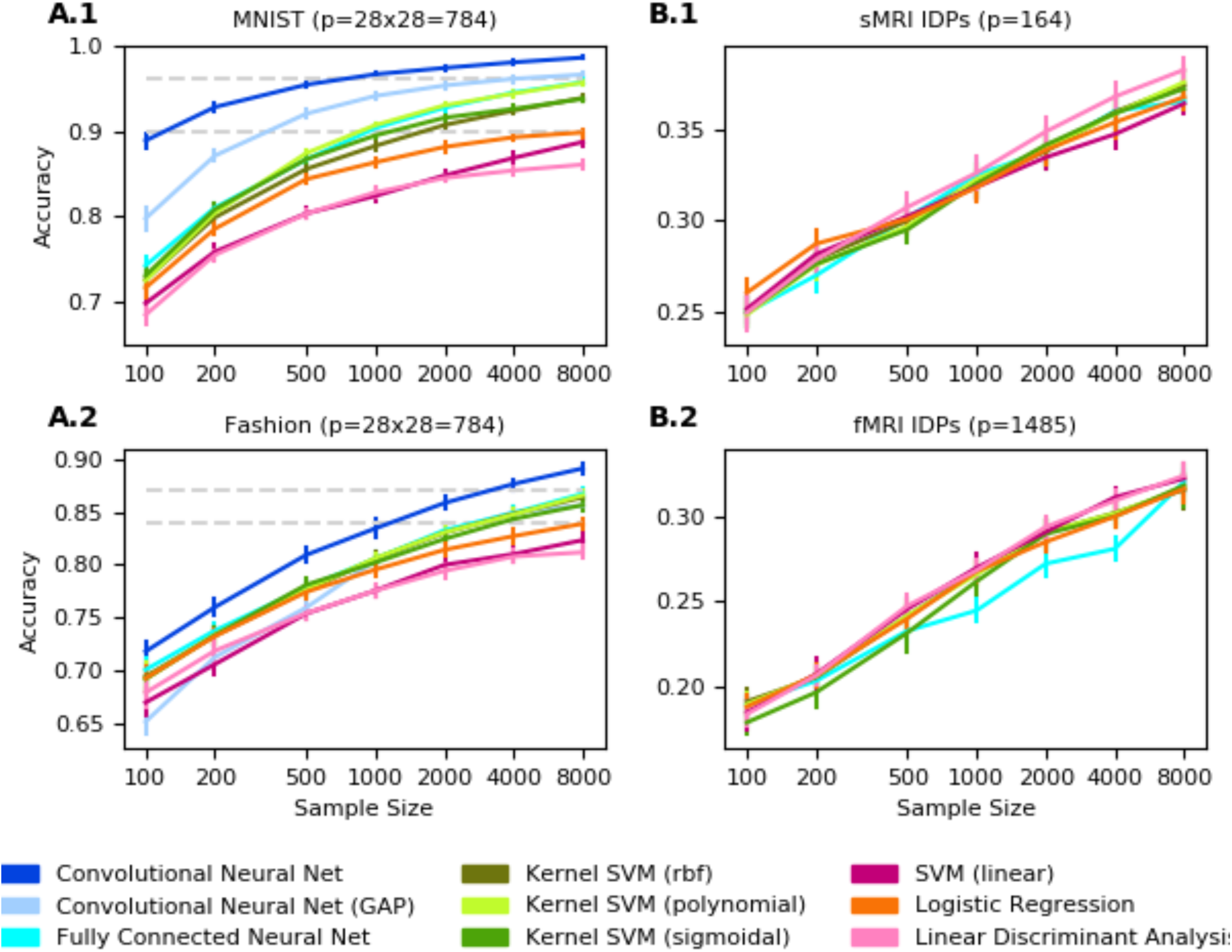
Classification performance increases with more powerful algorithms in two machine-learning reference datasets, but not in brain-imaging datasets. Shows performance scaling of prediction accuracy (y axis) as a function of increasing sample size (x axis) for generic linear models (red tones), kernel models (green tones), and deep neural network models (blue tones). All model performances are evaluated on the same independent test set to obtain comparable out-of-sample estimates of prediction accuracy that approximate external validation. A.1) In handwritten digit classification on the MNIST dataset, the classes of 3 linear, 3 kernel, and 3 deep models show distinct scaling behavior: linear models are outperformed by kernel models, which are, in turn, outperformed by deep models. The prediction accuracies of most models start to exhibit saturation at high sample sizes, with convolutional neural network models approaching near perfect classification of 10 digits. A.2) As a more difficult successor of MNIST, the Fashion dataset is about classifying 10 categories of clothing in photos. Similar to MNIST, linear models are outperformed by kernel models, which are outperformed by deep models, with expectedly smaller prediction accuracies. In contrast to MNIST, the performances of the model classes are harder to distinguish for low sample sizes and begin to fan out with growing sample size. The differences in prediction performance are less pronounced, and accuracies saturate more slowly with increasing sample size. In the Fashion dataset, more images are necessary for kernel and deep models to effectively exploit nonlinear structure to supersede linear models. B) Image-derived brain phenotypes (IDPs) provided by the UK Biobank were used to classify subjects into 10 subject groups divided by sex and age. The number of categories is equivalent to and the feature number p is similar to MNIST and Fashion. In both commonly acquired structural (sMRI) and functional (fMRI) brain images, linear, kernel, and deep models are virtually indistinguishable across all examined training image sets, and prediction accuracies do not visibly saturate. To the extent that complex nonlinear structure exists in these types of brain images, our results suggest that this information cannot be directly exploited based on available sample sizes.

Next, we examined more closely how the difference in 10-class prediction problem scales with increasing sample sizes in the MNIST and Fashion datasets. The sample size necessary to saturate the prediction performance of a classifier relates to the complexity of patterns that can be reliably derived from amount and dimensionality of observations available for model training. The out-of-sample prediction performance of both linear and kernel models saturated with increasing sample size in MNIST. Performance of linear models (logistic regression) grew by 14.67 p.p. from 71.64±2.62% to 86.31±1.06 % when the data availability grew from 100 to 1,000 observation, while the performance for shallow nonlinear models (i.e., RBF-SVM) grew by 15.60 p.p. from 72.58±2.34 % to 88.23±1.11 %. Improvements reduced to 3.50 p.p. (5.62 p.p. for RBF-SVM) from 1,000 to 8,000 image observations. This saturation of prediction accuracy was less pronounced when applying the same learning algorithms to images of the Fashion dataset: Linear models (logistic regression) grew by 10.22 p.p. from 69.34±2.95 % to 79.55±1.67 % from 100 to 1,000 observations, but only 4.31 p.p. from 1,000 to 8,000 observations. Shallow nonlinear models (SVM-RBF) grew by 11.08 p.p. from 69.50±2.89 % to 80.58±1.57 % from 100 to 1,000 observations, but only by 5.78 p.p. from 1,000 to 8,000 observations.

In sum, consistent with our expectations, nonlinear methods like kernel SVMs progressively outperformed linear models like logistic regression as the number of observations available for model building grew larger. Additionally, the prediction performance slowly saturated when increasing the number of observations beyond 1,000 example images. Both effects were more pronounced in the simpler prediction goal of digit classification (MNIST) than in the more complex clothing classification (Fashion). That is, in the more challenging prediction problem, the richer nonlinear models could learn even more as the amount of available image data grew. More complex classification tasks needed a higher number of samples to saturate the models. Our empirical scaling results confirm that these standard machine learning datasets contain nonlinear structure that our nonlinear kernel models can take advantage of with large numbers of observations.

### Model performance on brain-imaging data scales similar to linear models

To assess how the performance of common linear models and nonlinear kernel models behaves on brain-imaging data, we created a range of classification scenarios using a currently largest brain-imaging dataset - the UK Biobank Imaging. We tried to specifically design the classification problems to be similar to key properties of MNIST and Fashion to facilitate comparison of the results between brain-imaging and machine learning reference datasets. As a classification target, we constructed a 10-class target variable based on subjects’ age and sex in analogy to the data shape in MNIST and Fashion. Our UK Biobank data provided structural and functional MRI scanned in 9,300 subjects. We evaluated different views of the underlying brain data, corresponding to distinct and often used forms of brain-imaging data analysis: UK Biobank provides high level ‘imaging-derived-phenotypes’ (IDPs) of resting state functional brain MRI (ICA-based functional connectivity), and structural brain MRI (regional gray and white matter volumes), which brought the >100,000 grey-matter voxels per brain scan to a dimensionality that is akin to the dimensionality of MNIST and Fashion: fMRI p=1,485 variables, and sMRI p=164 variables. Additionally, to explore data analysis scenarios focused on voxel-level statistics (e.g., general linear model such as implemented in SPM and other common brain-imaging analysis software packages) we reduce the raw sMRI voxels to MNISTs dimensionality of 784 variables by different feature selection and dimensionality reduction techniques. Finally, in preparation of our analysis of modern convolutional neural networks, we extract central 2D sMRI slices for each anatomical plane.

Although we have made an effort of designing brain-imaging classification problems with similar properties to MNIST and Fashion in terms of number of classes, features, and sample size, we recognize that the brain-imaging classification problems are still more complex (e.g. age relates to the brain features in a continuous way and people’s brain age differently so there might be high overlap between the classes). We therefore ran additional control analyses on binary (sex) classification and continous (age) regression (Fig. 4).

**Fig. 3:**
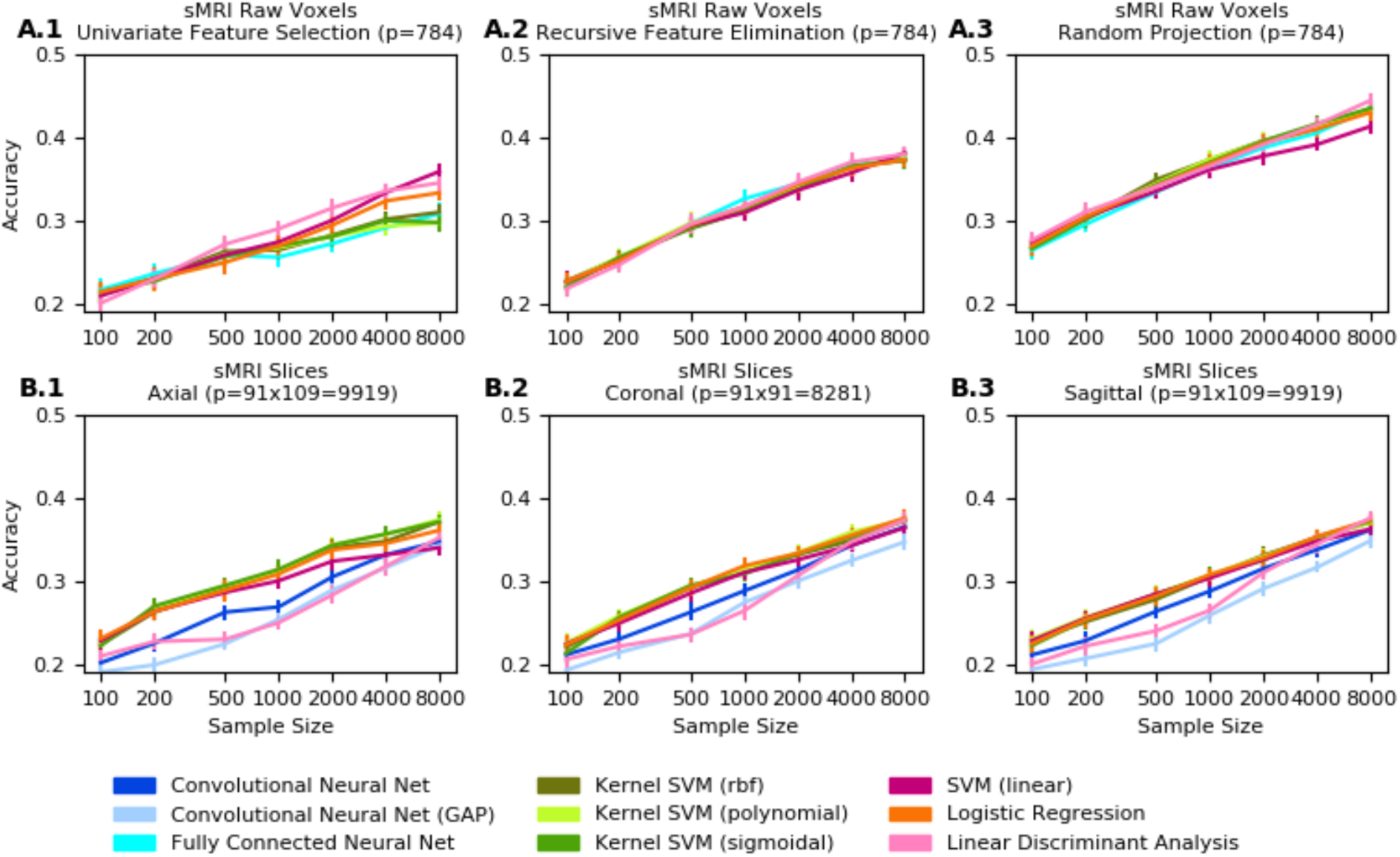
More powerful predictive algorithms fail to improve classification accuracy for several representations of structural brain images. A) Whole-brain grey-matter information from structural brain images (sMRI, UK Biobank) was summarized by three widely used feature selection/engineering techniques. The original tens of thousands of grey-matter voxel volumes were reduced to 784 variables for comparability to the dimensionality of MNIST and Fashion (Fig. 1), as a basis for learning predictive patterns for classifying 10 sex/age groups. Prediction accuracies achieved using univariate feature selection (relevance tests independent for each variable) are outperformed by recursive feature elimination (accounting for conditional effects between variables), which in turn is superseded by random projections (low-rank transformations of all original variables). None of these dimensionality-reduced brain images lead to systematic differences between linear (red tones), kernel (green tones), and deep (blue tones) model classes. In particular, fully connected deep neural networks did not visibly outperform the examined classical linear or kernel methods. B) Axial, coronal, and sagittal slices of whole-brain grey-matter images (sMRI) were used for 10-group categorization. In contrast to classifying 2D images of digits and clothing into 10 categories (Fig. 2, A.1-A.2), classifying 2D images of brain anatomy into 10 age/sex groups does not exhibits obvious performance differences between linear, kernel, and deep models. These analyses again indicate scarcity of easily exploitable nonlinear structure in common sMRI brain scans for present sample sizes.

**Fig. 4:**
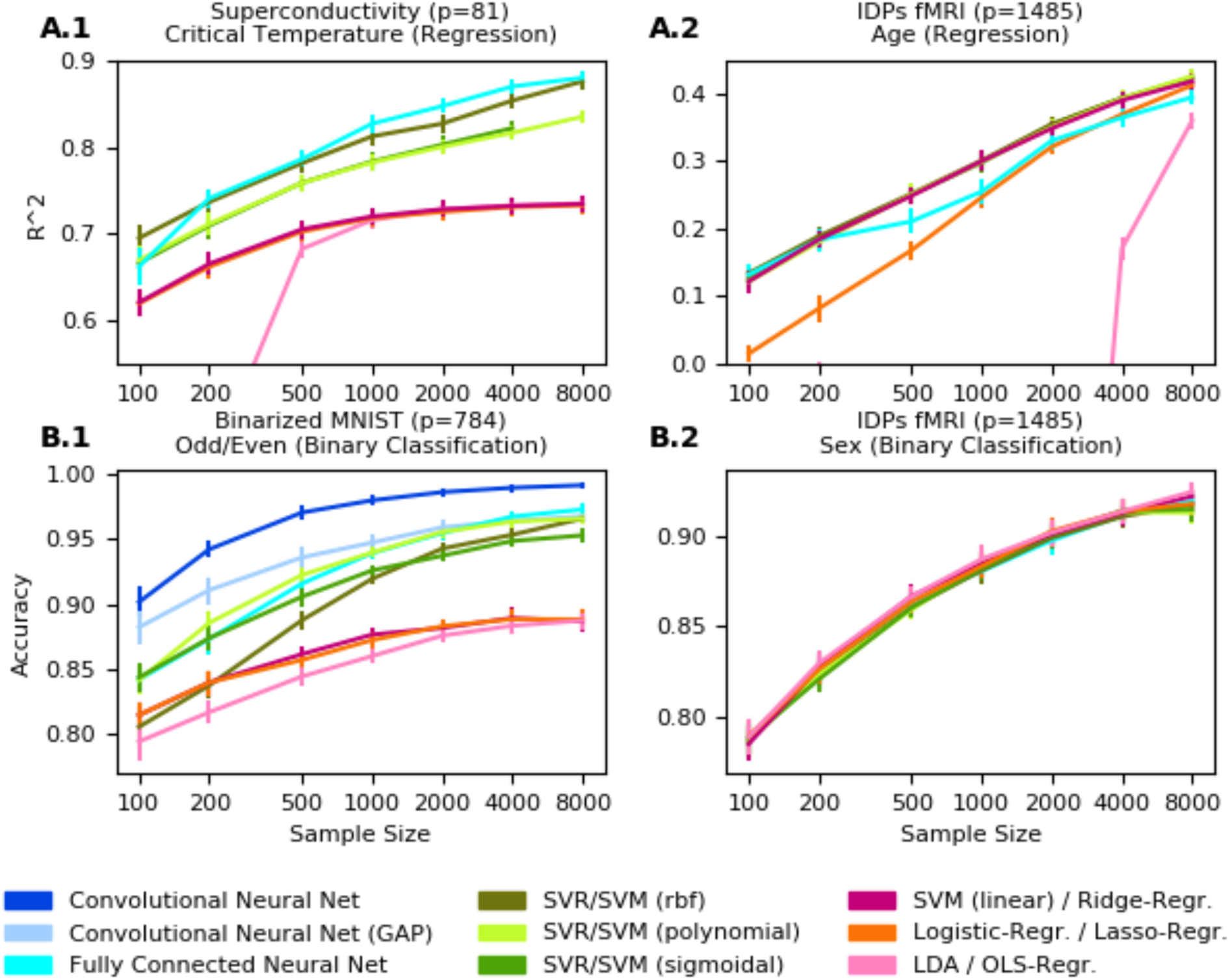
Quantitative and qualitative prediction shows exploitable nonlinearity in different open datasets, but not in functional brain images. To complement our classification of 10 age/sex-stratified groups (Fig. 2-3), we decoupled the stratified prediction goal by separately examining continuous age regression and categorical sex classification using linear (red tones), kernel (green tones), and deep (blue tones) models. In the superconductivity benchmark dataset (A.1), critical temperature was predicted based on 82 physical properties like thermal conductivity, atomic radius, and atomic mass (Hamidieh 2018). Here kernel and deep models clearly outperform linear models as measured by out-of-sample explained variance (R^2^). In contrast, in age prediction based on functional brain scans (A.2), the best performing linear model (Ridge regression) has scaling behavior virtually identical to kernel and deep models. This age prediction using fMRI scans is conceptually similar to previous analyses on brain maturity that reported 55% explained variance on 238 subjects aged 7-30 years (Dosenbach et al. 2010). These investigators noted “asymptotic maturation toward a predicted population mean maximum brain age of ∼22 years […] The fitted models mainly differed in their predictions for younger ages.” In our much older UK Biobank participants (62.00+/-7.50 years) we reach ∼40% explained variance, while the learning curves suggest further performance gains as more brain data become available. We identify a similar discrepancy between machine learning and brain-imaging datasets in the binary classification setting. In even (0, 2, 4, …) vs. odd (1, 3, 5, …) digit classification on MNIST (B.1), kernel and deep models diverge from linear models in classification accuracy as the sample size increases. However, the kernel and deep models are not superior in sex classification based on fMRI data (B.2), where all examined models showed virtually identical prediction performance.

In our analyses of brain-imaging data (Fig. 2-B), we observed different scaling behavior compared to benchmark machine learning datasets for common linear models and nonlinear kernel methods. Across the different views of the brain scans, model performance converged on the same pattern: For each sample size, accuracies for all examined models were mostly statistically indistinguishable. In this example of the currently largest brain-imaging dataset, we did not observe signs of accuracy saturation. That is, doubling the sample size yielded stable, mostly indistinguishable gains in accuracy. For example for sMRI IDPs (Fig. 2-B.1) at a full training set of 8,000 observations, the best kernel model (polynomial-SVM: 37.58±2.08 %) performed indistinguishably from the worst linear model (LDA: 38.26±1.46 %). Performance of linear models (logistic regression) grew by 5.78 p.p. from 26.02±1.98 % to 31.80±1.92 % (7.23 p.p. from 24.92±1.81 % to 32.14±2.44 % for RBF-SVM) from 100 to 1,000 observations, indistinguishable from the 4.97 p.p. (5.29 p.p. for RBF-SVM) for 1,000 to 8,000 observations. Qualitatively equivalent results were observed for different dimensionality reduction techniques (Fig. 3-A) in regression and binary classification settings (Fig. 4), and, importantly, for different prediction targets - sex, age, and the combined 10-class variable (Fig. 2+4). In contrast to the examined machine learning reference datasets, brain-imaging data revealed neither saturation of prediction accuracy with increasing sample size nor performance gains from kernel models with higher expressive capacity.

### Performance of deep neural network models

Finally, we compared the performance of deep neural network models on our brain-imaging data. Both fully connected neural network architectures and common variants of convolutional neural networks systematically differed in the performance on the machine learning benchmark datasets compared to accuracy profiles in brain-imaging data.

Fully connected neural networks outperformed all linear models and all but the best performing kernel model on the MNIST and Fashion datasets (Fig. 2-A), but not for any examined representation of brain imaging data. Regarding sMRI and fMRI IDPs (Fig. 2-B), sMRI voxels (Fig. 3-A) and whole sMRI slices (Fig. 3-B), fully connected neural networks performed indistinguishably from both kernel and linear models. The same held true for our control analyses of the regression and the binary classification (Fig. 4).

Convolutional neural networks showed analogous scaling behavior. On both MNIST and Fashion datasets, convolutional neural networks outperformed the best performing kernel model by 2.93 p.p. from 95.63±0.79 % to 98.56±0.36 % and by 3.44 p.p. from 85.63±1.32 % to 89.07±1.20 % respectively at 8,000 observations (Fig. 2-A). In contrast, on sMRI slices convolutional neural networks outperformed neither linear nor kernel models (Fig. 3-B). Central axial, coronal, and sagittal slices yielded comparable prediction accuracy for logistic regression (36.09±1.38 %, 37.58±1.81, 37.35±1.21 % respectively) and convolutional neural networks (34.74±1.65 %, 36.52±1.83 %, 36.15±1.82 % respectively). Thus, machine learning reference datasets and brain-imaging data differed in that convolutional neural networks excel on the former, but fail to improve over simpler methods on the latter.

Taken together, we described three important ways in which predictive models performed systematically differently between the machine learning reference datasets MNIST and Fashion on the one hand and brain-imaging data on the other hand. First, performance of linear models saturated with increasing sample size on MNIST and Fashion (less so kernel and deep models), but not on brain-imaging data. Second, kernel models consistently outperformed linear models in MNIST and Fashion, which was not the case in our analyses of brain-imaging data. Third, deep models clearly outperformed linear and kernel models on MNIST and Fashion, but not on our brain-imaging data. Our quantitative investigation can therefore be taken to argue that the largest brain-imaging dataset currently at our disposal still do not reliably enable exploitation of complex configurations in data by modern pattern-learning algorithms.

## Discussion

Do today’s kernel methods or deep neural network algorithms provide advantages for analyzing common brain images by drawing on shallow-nonlinear or deep-nonlinear information? Can the triumphs of kernel and deep methods in machine learning datasets be obtained in currently available brain-imaging datasets? Our study initiates quantitative answers to these increasingly important questions. We have profiled the degree to which complex nonlinear relationships can be extracted and exploited to improve prediction performance over linear models in structural and functional brain data from the UK Biobank. Our careful analyses delineate the scaling behavior of prediction performance with increasing number of training observations for three classes of learning algorithms: linear, kernel, and deep models. Importantly, we here replicated performance gains escalating to nonlinear and then to hierarchical nonlinear models reported in performance benchmarks from machine learning. Such improvements with increasing model complexity were not consistently apparent in the currently largest population brain-imaging cohort.

### Present sample sizes are too small to even fully exploit linear modeling

A key finding that emerged from our analyses pertains to the performance trajectory of linear models with gradually growing sample size (i.e., empirical sample complexity). Our prediction outcomes from brain-imaging data and reference datasets from machine learning differed in the way that the prediction accuracy increased with the availability of data from additional observations. In the machine learning datasets, we expected, and indeed observed, saturation of prediction performance of linear models. This effect was most prominent when carrying out digit classification on MNIST by means of logistic regression and linear discriminant analysis: The prediction performance increased rapidly as we grew the sample size from 100 to 1000 available images. Subsequently, the prediction accuracy approached a plateau given that we noticed hardly any additional improvement in prediction accuracy when doubling the sample size from 4,000 to 8,000 example images.

In contrast, in our analyses of brain scans, this step up in the number of images available for model building yielded continuous gains in prediction performance for all examined brain-imaging modalities and data representations. Across different MRI protocols to measure brain tissue, we determined that increasing the number of participants from 4,000 onward entailed steady performance gains in prediction accuracy. Importantly, the size of the largest available brain-imaging repositories was insufficient to saturate the learning capacity of even simple linear models, which were here applied in a “multivariate” fashion pooling brain information from many parts of the brain (Haynes 2015; Woo et al. 2017). Our findings suggest that, approaching brain scans obtained from up to 10,000 participants, the prediction capacity of common linear models is not yet fully exhausted.

We deem the uncovered prediction reserve for the linear modeling regime important in at least two ways. First, this scaling behavior provides new arguments for the common criticism that there may be limited information contained in brain-imaging data like MRI that can be usefully exploited for prediction in real-world applications (Woo et al. 2017). For instance, in the case of functional MRI, blood flow and oxygenation do not provide an immediate read out neuronal activity, and operate at coarser-grained temporal and spatial scales than actual electrochemical information processing in neuron populations. Some authors have claimed that “fMRI is as distant as the galvanic skin response or pulse rate from cognitive processes” (Uttal 2011). Despite these caveats, there have been a number of encouraging findings. For instance, functional brain connectivity could be shown to provide a neural fingerprint to make accurate predictions of cognitive performance (van den Heuvel et al. 2009; Pamplona et al. 2015; Finn et al. 2017). Yet, such promising reports are controversially discussed in the community, and have been judged hard to replicate by some investigators (Kruschwitz et al. 2018).

Our results favor the more optimistic interpretation scenario and suggest that we are not yet fully exploiting predictively useful information in brain-imaging data. Not even simple linear models have reached plateaus of prediction performance in our sex and age predictions from common MRI measurements at present sample sizes. As such, we are likely to be far from reaching the limits of single-subject prediction accuracy by leveraging brain-imaging data.

Our finding of unexhausted linear modeling reserve may have considerable ramifications. This is because systematic evaluations of modern machine learning in brain-imaging (He, Kong, et al. 2018; Vieira et al. 2017; Plis et al. 2014) as well as a large amount of studies applying complex nonlinear models in brain-imaging often have operated under the implicit assumption that linear effects are already sufficiently characterized with their prediction scaling as sample size increases. Typically, carefully characterizing linear effects provides a solid basis to compare against more complex nonlinear models. Many elaborate machine learning methods can be viewed as extensions of classical linear regression (J. Friedman et al. 2001). If there is insufficient data to estimate the parameters of a simple linear model, then it is even less likely that the parameters of even more data-hungry nonlinear models can be estimated with satisfaction. The recommendation emerging from the present investigation is that (regularized) linear predictive models are likely to serve as a formidable starting point for single-subject prediction in larger future brain-imaging datasets in health, and potentially also disease, for the foreseeable future.

### Limited success of exploiting nonlinear information in currently available brain-imaging data

If nonlinear structure exists in a given dataset and is exploitable for a specific prediction goal and sample size, we expect an increase in prediction performance as we switch from classical linear models to always more expressive models (Jerome Friedman et al. 2001; Ghahramani 2015; Bzdok & Yeo 2017). In a second set of analyses, we confirmed the expected increase in prediction performance with growing capacity to represent complicated patterns in one of the most popular machine learning benchmark datasets (digit image classification in MNIST), as well as a more difficult companion dataset (clothing image classification in Fashion). In particular, kernel methods consistently outperformed commonly used linear models, with accuracy gains of 4.03 and 1.76 p.p. on MNIST and Fashion, respectively, on average.

Our ability to quantify the detection of nonlinear data configurations in MNIST and Fashion to distinguish 10 categories of images (10 digits or types of clothing) corroborates our application of kernel SVMs as a viable and effective tool to test for the existence of predictive nonlinear patterns in data from the brain-imaging domain. When information carried in these brain measurements are related to the prediction goal in complicated ways, an SVM with a nonlinear kernel extension is expected to largely outperform linear models if provided with brain scans from enough participants.

However, we did not observe systematic performance gains in prediction accuracy when examining brain images from UK Biobank, although we empirically replicated them in our analyses on the MNIST and Fashion datasets. We found that none of the three nonlinear kernel models clearly outperformed linear models on structural or functional brain scans. This finding is particularly apparent in the fMRI data widely used as for computing functional connectivity between brain regions and networks, where all examined models perform nearly identically across the sample sizes empirically simulated in our study. In fact, kernel and linear methods performed virtually indistinguishably in a wide range of analyses of brain-imaging data from the currently largest biomedical dataset - the UK Biobank - designed to be approximately representative of the UK general population (Fry et al. 2017). A similar conclusion on fMRI data has been drawn years ago by Cox and colleagues (2003). These investigators noted that “in spite of many conceivable sources of nonlinearity in neural signals, the nonlinear […] SVMs used did not significantly outperform their linear counterparts.”

Limited gains in prediction performance from adopting always more elaborate nonlinear models may turn out to be a common property of brain-imaging analysis settings with sample sizes in the order of thousands of subjects. We therefore examined different neurobiological measurements as obtained by 3T MRI scanning lending insight into brain anatomy and intrinsic functional coupling between brain regions and networks. We further assessed different representational windows into the brain scans, from the application of various data representation methods to hand crafted features to raw (1D) voxel data to feature engineering/selection approaches to whole-brain slices (2D). Moreover, we have considered different prediction goals - age, sex, and stratified age/sex groups - known to explain large amounts of variability in brain MRI data (Miller et al. 2016). Nevertheless, in our analyses of canonical machine learning datasets, but not in brain-imaging data, linear models were reliably outperformed by all examined nonlinear kernels. Even easy-to-predict phenotypes, such as age and sex, failed to show reliably improved prediction performance when switching to current nonlinear models for sample sizes in the order of thousands of subjects. Thus, we are enticed to speculate that even harder to define and trickier to measure concepts, such as IQ, social cognition capacity, and mental health diagnoses, will seldom achieve performance gains from deploying complex models at similar sample sizes.

Our study may be myopic regarding some conceivable conclusions. The fact that most analyses showed highly comparable prediction accuracies between linear models and models with kernel extensions in brain images allows for several possible interpretations. On the one hand, it may be the case that there exist few salient nonlinear relationships in the examined types of brain data. In this case, linear models would be expected to be more efficient and effective at extracting the crucial patterns that are instrumental for the prediction goal. On the other hand, it may be the case that nonlinear configurations truly existing in the brain data cannot be easily used to serve our prediction goals given the size of currently available brain-imaging repositories. The nonlinear interactions in brain-imaging data could be so intricate that it would take a substantially larger sample size to reliably capture and leverage them from the brain data (Gelman & Hill 2007). Additionally, the lens through which the here charted nonlinear methods “see” principled patterns in the data may not be aligned well with the kind of nonlinearity present in these types of brain scans (i.e., mismatch of *inductive bias*). As a limitation of the present investigation, we cannot provide a definitive answer as to which of these possibilities is more pertinent.

As a practical consequence, the scarcity of exploitable nonlinear structure in our brain-imaging data suggests that linear models will continue to play a central role in analyzing brain scans like MRI measurements, at least over the next few years (He, Kong, et al. 2018; Bzdok & Ioannidis 2019; Smith & Nichols 2018). The additional representational expressivity that modern nonlinear models provide comes at the cost of an increased risk of overfitting (Varoquaux et al. 2017) and typically added challenges in interpretability of the estimated models (Bzdok et al. 2019; Lundberg & Lee 2017; Chen et al. 2018). Our empirical results suggest that, for sample sizes available today, the added costs of implementing one of the current more complex models in computational requirements and technical knowledge may rarely be justified by the theoretical potential for better prediction performance on regular MRI brain scans.

### Deep learning did not universally improve prediction performance in brain-imaging data

Finally, we consider our results on the value of current deep neural network algorithms in imaging neuroscience. Three trends appear to stand out in the existing brain-imaging literature. First, it is noteworthy that there is still a suspicious scarcity of published MRI or positron emission tomography (PET) studies that unambiguously demonstrate substantial gains from applying deep learning techniques. In line with this, Vieira and colleagues (2017) noted that “despite the success of [deep learning] in several scientific areas, the superiority of this analytical approach in neuroimaging is yet to be demonstrated”. In domains like computer vision or natural language processing, deep neural networks have already dramatically improved state-of-the-art prediction performances (Goodfellow et al. 2016; LeCun et al. 2015). However, in the application of neuroimaging data analysis, a similar revolution has not materialized for most common prediction goals, despite considerable research effort and few successful exceptions, such as for the goal of image segmentation (Choi & Jin 2016; Kamnitsas et al. 2017; Li et al. 2018; Kather et al. 2019) and image registration <u>(Balakrishnan et al. 2018; Yang et al. 2017)</u>.

Second, deep learning approaches have repeatedly been found to perform worse or indistinguishably well compared to simpler baseline models when predicting demographic or behavioral data (Cole et al. 2017; Jang et al. 2017; He, Kong, et al. 2018). For instance, Cole and colleagues (2017) showed that deep convolutional neural networks did not outperform Gaussian process models when predicting brain age from structural MRI data in ∼2,000 healthy participants. Consistently, He and colleagues (2018) found that three different deep neural network architectures did not outperform kernel regression at predicting a variety of phenotypes, including age, fluid intelligence, and pairs-matching performance from whole-brain connectivity profiles derived from functional MRI data. Moreover, in the 2019 ABCD challenge, kernel ridge regression outperformed deep learning approaches in predicting fluid intelligence from structural MRI (Mihalik et al. 2019), while the four best predictive modeling approaches did not utilize deep learning in the TADPOLE challenges to predict progression in Alzheimer’s disease (Marinescu et al. 2018). In general, even many reports of modest improvements from using deep neural networks are controversially discussed in the brain-imaging community.

Third, contrary to many scientists’ expectations, gains from machine learning algorithms with the capacity to represent complicated nonlinear relationships in brain-imaging appear to decrease or stagnate when incorporating data from more sources or acquisition sites. This recurring observation suggests overfitting or possible publication bias. The general expectation is that improving model building with additional training data should improve prediction performance of complex nonlinear models, especially when moving from dozens or few hundreds to thousands of participants. In contradiction with this intuition, Arbabshirani and colleagues (2017) pointed out that “the reported overall accuracy decreases with sample size in most disorders”. Specifically, in the context of deep learning, Vieira and colleagues (2017) noted that “the pattern of difference in performance did not seem to vary systematically with sample size”. Woo and colleagues (2017) concluded that their “survey reveals evidence for such [publication] bias in predictive mapping studies”. These circumstances were speculated to reflect increasing heterogeneity of larger patient samples and inter-site differences in the process of data acquisition. However, these considerations also cast some doubt on the high expectations about embracing deep neural network applications to the types and amount of brain-imaging data that is available today.

Our results from careful model profiling are in line with these previous reports and observations from imaging neuroscience studies. Certain deep neural network models consistently outperformed all kernel models and all linear models in our analyses of MNIST and Fashion datasets. We did not see such effects in our analyses of brain-imaging data. Here, the majority of models performed statistically indistinguishably for several investigated imaging modalities, data representations, and predicted target variables. This lack of consistency in performance differences between model classes - in the world’s largest biomedical dataset - is instructive as it lends credence to reports of the potential for overfitting or publication bias in the field.

In carefully controlled experiments, even null results can be ‘evidence of absence’ <u>(Hacking 2001; Thompson and Scurich 2018)</u>. However, due to the high flexibility of deep learning, it is nearly impossible to fully explore all possible combinations of hyperparameters and model architecture choices (Goodfellow et al. 2016, chap.11.4). Hyperparameter search is computationally expensive so that only a well-chosen subset of candidate hyperparameter combinations can reasonably be evaluated in any given study. Thus, negative results in training deep neural networks are often disregarded as they leave open the possibility of insufficient hyperparameter tuning. A common response to a negative result in deep learning is to challenge the range or granularity of the hyperparameter grid, or to question the model architecture or the data preprocessing choices. We are aware that the same criticism can be applied to our own negative results.

Previous work from the neuroimaging community (He, Kong, et al. 2018) has made similar claims in stating that “deep neural networks [do not] outperform kernel regression for functional connectivity prediction of behavior”. Yet, their experimental analysis setup may not be able to fully dismiss the critique of insufficient hyperparameter optimization in deep models. In this way, our work provides a critical addition to the existing literature, by providing some support to the idea that kernel models for exploiting nonlinearity might not even be expected to outperform simpler linear models. As such, hurdles in exploiting even hierarchical nonlinearity may have been anticipated before the deep learning era in brain-imaging.

While still an active field of research, two theoretical attempts are generally evoked to explain the exceptional performance of deep neural networks on a variety of applications; besides the mathematical proofs that deep models are able to approximate arbitrarily complex prediction rules given sufficient training observations (“Universal approximation theorem”, Csáji 2001). One view states that the hierarchical, compositional nature of deep learning allows for a particularly parsimonious representation of some forms of nonlinear structure embedded in data (Bengio et al. 2007; Mhaskar et al. 2017). In analogy to a world map, the globe can be seen as compositional of continents, which split into countries, which split into provinces, which split into cities, and so forth. This hierarchical structure may be efficiently represented by a multilayer nonlinear processing layers in artificial neural network models, where individual layers correspond to streets, districts, cities, etc. The same nested structure applies to human language, and thus written text and recorded speech data.

Based on this notion, one should expect deep neural networks to improve upon linear and kernel models in a given domain only if there exists exploitable hierarchical and nonlinear structure in the data. Compared to kernel methods, deep neural networks tend to have orders of magnitude more parameters and often require a correspondingly high number of training observations. The recent empirical successes of deep learning are largely attributed to the growing availability of extremely large datasets, especially in domains building on Internet-scale image and natural language data, as well as availability and affordability of computation (Goodfellow et al. 2016). Deep neural networks appear to violate theorems of statistical learning theory in that some highly overparameterized models still generalize unexpectedly well to new data (Zhang et al. 2016). Nevertheless, we generally expect kernel methods to require fewer training observations to extract useful nonlinear structure in data (Klambauer et al. 2017). Additionally, kernel methods have only a few hyperparameters, which allows for a more comprehensive search of the optimal model subspace for the data at hand. Given their typically more efficient empirical hyperparameter tuning, kernel methods thus have the practical potential to select model instances that are better suited to identify and exploit nonlinear interactions existing in the data in smaller sample sizes than deep neural networks. However, kernel methods do not have the ability to take advantage of complex hierarchies in representing patterns in the data effectively. Thus, kernel methods are, arguably, preferable when the data do not contain exploitable hierarchical structure.

The second commonly evoked explanation for why deep neural networks may outperform earlier methods is specific to convolutional deep neural networks. This type of network was designed specifically for processing natural images. Convolutional neural networks were motivated by the organization of the visual cortex, in that individual neurons respond to stimuli in only a limited receptive field and as such exploit the locality of information in images - the way in which neighboring pixels describe the same phenomenon. Similar to the visual cortex, convolutional neural networks consist of a hierarchy of layers of locally sensitive feature-detectors. In contrast to the visual cortex, each feature-detector is recycled - convolved - for each position in the image. This reuse of feature detectors at different positions leads to the inbuilt bias called translation invariance. For example, a feature-detector sensitive to a cat would work independently of where the cat is located in an image. This inductive bias for translation invariance introduces a useful and domain-compatible simplification in the analysis of natural images because translation invariance is an important property of the physical world - a cat remains a cat independently of where in space it is located.

However, in contrast to MNIST, Fashion, and most other common computer vision datasets, the brain has a naturally meaningful topography in an MRI image. The scanned individual’s head position in the scanner and the subsequent mapping to standard atlas space is widely established. For a majority of the possible prediction goals, there is no need for a translation invariant feature detector to search the whole volume for a particular representational phenomenon - any given brain region is located in a fixed predefined space. It is therefore unlikely that using convolutional neural networks invariably improves prediction performance on standard-resolution brain-images in common reference space.

Importantly, the take home message from our present work is *not* that deep neural networks will never be a useful tool for brain-imaging data. Deep convolutional neural networks should be particularly useful when confronted with anomalies that can be anywhere in the brain, such as for detecting tumors or quantifying abnormal lesions in multiple sclerosis (Pereira et al. 2016; Havaei et al. 2017; Kamnitsas et al. 2017; Valverde et al. 2017). Nevertheless, our results suggest that for a variety of brain-imaging applications, there might be little advantage to using current convolutional deep neural network models for answering neuroscientific research questions at today’s sample sizes.

## Material and Methods

### Three reference datasets: Three test beds for interrogating the scaling behavior of predictive models

#### MNIST

This dataset (Modified National Institute of Standards and Technology dataset; LeCun & Cortes 2010) may be the most used reference dataset for research and development in the machine learning community. Its properties are well understood because a large number of models have been benchmarked on MNIST. This classical dataset provided a convenient starting point for the present study, allowing us to reproduce and quantitatively characterize different properties of machine learning algorithms in a controlled setting. MNIST provides 70,000 images of handwritten digits (‘0’-’9’) to be classified based on the raw pixel information. Each of these grayscale images, consists of 28×28 pixels, that is, 784 intensity values in total per digit image.

#### Fashion

To quantitatively characterize the effects of more complex data patterns with a more challenging prediction goal, we also analyzed our predictive models on Zalando’s recent Fashion dataset <u>(…Fashion-MNIST", Xiao et al. 2017)</u>. Instead of handwriting, the dataset consists of grayscale images of fashion products. Images are to be classified into 10 types of clothing (e.g., t-shirt, sweater, dress). MNIST has sometimes been found to be too easy to predict for very recent machine learning methods. The Fashion dataset was created with the intention to provide a more difficult pattern recognition problem than MNIST, while preserving the same number of classes (10 clothing categories), feature dimensionality (784 pixel intensities), and sample size (70,000 images). This conveniently allowed for using the same model architectures on both machine learning benchmark datasets and facilitated comparisons of model performance scaling.

#### UK Biobank

Our goal was to compare the known properties of MNIST and Fashion datasets to common types of brain images acquired in humans (rather than an exhaustive benchmarking of deep learning in neuroimaging in general). Such direct juxtaposition allowed identifying settings where model behavior extends from machine-learning datasets to brain-imaging datasets. Additionally, these analyses allow developing some first intuitions of the settings where extrapolation of effects should not be expected. UK Biobank imaging was a natural choice for the motivation behind our experiments (Sudlow et al. 2015). This resource is the largest existing biomedical dataset to date (Sudlow et al. 2015). Our data request of the UK Biobank brain-imaging initiative provided structural and functional magnetic resonance imaging (MRI) data for ∼10,000 subjects from the same scanning site (UK Biobank application number 25163, information on the consent procedure can be found at biobank.ctsu.ox.ac.uk/crystal/field.cgi?id=200). We centered our analysis on a complete set of UK Biobank individuals who simultaneously provided all the imaging modality of brain structure (sMRI) and function (fMRI), which resulted in a total sample size of 9,300 data points from brain images.

From the UK Biobank release, we compiled a multifaceted set of brain-imaging data, representing different modalities, as well as different views on a given modality. We derived 4 working datasets based on the brain images: a) Intrinsic neural activity fluctuations measured by functional MRI data, with pre-computed independent component analysis yielding 100 spatiotemporally coherent neural activity patterns, resulting in a feature space of 1,485 connectivity strengths between cleaned “network” components estimated using partial correlation. b) 164 atlas-derived features describing structural (T1-weighted MRI) MRI grey- and white-matter region and fiber tract summary statistics. For both region volume and fiber bundle microstructure estimates, the biologically meaningful features were provided directly by UK Biobank as “imaging-derived-phenotypes” <u>(Miller et al. 2016)</u>. c) ∼70,000 raw T1 voxel intensities, after gray-matter masking. d) Finally, axial, sagittal, and coronal T1 slices at the origin with resolutions of 91×109, 91×91, 91×109 voxels, respectively. Further details on data acquisition, data preprocessing, and IDPs are provided elsewhere (Miller et al. 2016; Alfaro-Almagro et al. 2018).

For the sake of comparability with the MNIST and Fashion datasets from machine learning, we generated a prediction target of 10 classes from all possible combinations of 2 sexes and 5 age quintiles. That is, the UK Biobank participants were divided into 5 male subgroups with four age cut-offs between 40-70 years and 5 female subgroups with these four age cut-offs. As our brain-imaging dataset provided a total of 9,300 individuals with multi-modal imaging information, at the time of our analysis, we randomly subsampled the much larger MNIST and Fashion datasets to the same sample size in our analyses.

Although we have designed brain-imaging classification problems with similar number of classes, features, and sample size as the MNIST and Fashion datasets, the brain-imaging classification tasks are still expected to be much more complex than the classification problems in MNIST and Fashion.

### Preprocessing procedures

In all analysis scenarios, the input variables were standardized by scaling variance to one and centering to zero mean across observations (Kuhn & Johnson 2013). In the special case of convolutional deep neural network models (cf. below), the images were standardized on the aggregate pixel statistics to keep all pixels on the same scale (He, Zhang, et al. 2018).

For sMRI images, we applied an additional dimensionality reduction step for the purpose of comparability with MNIST and Fashion. To keep the model complexity comparable between datasets (related to the input variables and thus the number of model parameters to be fitted when building a prediction model), sMRI data were reduced to 784 features, like MNIST, by three dimensionality-reduction methods. Each of these represents a popular, but distinct approach to feature selection/engineering <u>(Friedman et al. 2001; Goodfellow et al. 2016; Kuhn and Johnson 2013)</u>: a) Univariate feature selection (F-tests) selected input variables under the assumption of statistical independence between variables; b) recursive feature elimination was used in 4 steps of logistic regression, each discarding the 25% least predictive input variables of the current active variable set in each step (Guyon et al. 2002). This dimensionality-reduction strategy selects features under joint consideration of the current set of input variables; c) Gaussian random projections were used to re-express the original input variables in a set of latent factors that rely on underlying low-rank structure across the whole set of input variables (Bingham & Mannila 2001).

### Linear models: Imposing simple additive structure on the data

Three classes of models - linear, kernel, and deep - were used to comprehensively evaluate the prediction performance on each dataset (i.e., MNIST, Fashion, UK Biobank). Each level of model complexity represents the state-of-the-art of a key epoch in data analysis (Efron & Hastie 2016): Regularized linear models to handle large numbers of input variables (∼1990-2000), kernel support vector machines became go-to methods for many applications in bioinformatics and beyond from late 1990s (∼2000-2010), and deep neural network algorithms became an exponential technology very recently (∼2010-2019). Within each of these model classes, we used three commonly employed representative models, attempting to cover the range of approaches within each particular modeling class.

For the class of linear methods, we have selected a) linear discriminant analysis, b) logistic regression, and c) linear support vector machines (SVMs; without kernel extensions). Linear discriminant analysis is a popular generative classifier that finds linear combinations of features that best separates classes among the observations, which is closely related to other covariance-based analysis methods like PCA (Fisher 1936; McLachlan 2005). L2-regularized logistic regression (Cramer 2002) and linear SVMs (Cortes & Vapnik 1995) are commonly used discriminative models, and represent frequently chosen instances of generalized linear models and linear maximum-margin classifiers, respectively (Hastie et al., 2001). Penalty terms for regularization constraints were tuned by grid search.

### Nonlinear models: Feature enrichment by kernelization

In practice, many classification problems are not linearly separable in the original variable space. The classes to be distinguished can become separable after nonlinear transformation of the input data into a representationally richer high-dimensional space. Solving a prediction problem in this high-dimensional representation tends to be computationally prohibitive and became more tractable in practice by means of the kernel methods.

Kernel methods (Schölkopf et al. 2002) are able to efficiently map to high-dimensional input spaces by never explicitly computing coordinates of all data points in this enriched space, but relying on the pairwise similarities between observations instead. Many linear methods can be extended to the nonlinear regime by applying this so-called ‘kernel trick’. Arguably the most popular kernel estimators are kernel SVM variants (Schölkopf et al. 2002). These feature embedding extensions are still pervasively used today in many application domains because of their well-understood theoretical properties, robust estimation, and often competitive real world performance. The class of kernel approaches was, and often remains the go-to choice to identify and use complex nonlinear interactions that exist in the data.

We therefore decided to evaluate kernel SVMs with three of the most commonly used kernel types: a) The radial-basis-function kernel, which maps the data to bell curves centered around observations, and is popular partly because of its universal approximation capacity (Wang et al. 2004). b) The polynomial kernel, which maps to polynomial expansions of the original variables, which made these variable enrichments popular in text processing using natural language processing as it explicitly takes into account combinations of features (Gliozzo & Strapparava 2009). c) The sigmoidal kernel, which maps to the hyperbolic tangent - analogous to the “activation function” that introduce nonlinearity in the units of deep neural network architectures (Goodfellow et al. 2016) - and is popular due to its and relation to the shallow “perceptron” artificial neural networks (Lin & Lin 2003; Collobert & Bengio 2004). These three instances of kernelization for nonlinear enrichment of input variable information are probably the most frequently, and the ones implemented in the dominant software packages (libSVM, SVMlight, scikit-learn).

### Hierarchically nonlinear models: Nested nonlinear transformations in deep learning

In an increasing number of application domains, single nonlinear enlargement of the input variables have been superseded by deep neural network (DNN) models. As an extension of linear and kernel methods, DNNs can extract and represent even more complex patterns in data using an automatically derived hierarchy of nonlinear operations on the set of input variables (Goodfellow et al. 2016). This nested design principle allows the learning architectures to pick up on progressively more abstract intermediate representations from the data themselves - a form of “automatic feature engineering” that was static in kernel methods (Goodfellow et al. 2016). Even though more exotic artificial neural network architectures exist, fully connected deep neural networks and the more recent convolutional deep neural networks are among the most often employed types.

Analogous to our analyses based on linear and kernel models, we evaluated several common deep neural network architectures. a) Fully connected neural network algorithms: The input layer has p unit for p-dimensional input data. The input layer was followed by two fully connected layers of 800 units each, with rectified linear unit (ReLU) nonlinearities and 50% dropout probability. These consecutive nonlinear operations are followed by a final fully connected layer of 10 units, followed by a softmax output function corresponding to predicting the probability of the target classes.

b) For the small (p=28×28) MNIST and Fashion images, convolutional DNNs consisting of two sets of convolutional and max-pooling layers (with 16 and 32 filters of 3×3 pixels respectively, a 2 pixel pooling size, and ReLU nonlinearities), followed by a fully connected layer of 128 units with ReLU nonlinearities, and a final fully connected layer of 10 units with softmax output corresponding to the target classes. Most tutorials and examples (e.g., for TensorFlow and PyTorch) use variations of these two basic architectures.

c) For completeness, we implement a third architecture in which the final max-pooling operation is replaced by global average pooling (GAP) - a popular approach to avoid overfitting by reducing the total number of model parameters <u>(Lin et al.; He et al. 2016)</u>.

For the larger (p=∼100x∼100) brain image slices, we inserted two extra convolutional and max-pooling layers to achieve a sufficient reduction in dimensionality before connecting to the prediction-generating output layer.

The parameters of the deep models were trained with the ADAM optimization algorithm (Kingma & Ba 2014): 160 epochs, with 500 gradient updates per epoch, a batch size of 32, learning rate reduction by 0.5 after 3 epochs without improvement, early stopping after 10 epochs without improvement. All deep models were trained with training-set-dependent levels of L2 regularisation as the only hyperparameter.

It is important to note that the goal of our study was not to benchmark highly specialized DNNs architectures from the recent neuroimaging literature. Instead, we decided to rely on established best practices to implement simple, straight-forward architectures as representatives for commonly used deep learning approaches.

### Model selection and evaluation: Hyperparameter tuning and cross-validated out-of-sample prediction

To estimate the prediction accuracy that we expect to obtain in new observations sampled from the population, cross-validation was computed as a gold-standard to obtain out-of-sample accuracies (J. Friedman et al. 2001). We repeatedly splitted the observations, sometimes called “monte-carlo cross-validation”, into a training set, as well as a validation set used for model selection (i.e., hyperparameter choice) and a test set used for model evaluation with 650 observations each in the sample complexity analyses (cf. below). In the UK Biobank, we considered 9,300 subjects with the brain-imaging data of interest. This random splitting was repeated 20 times. Training, validation, and test set were drawn exactly once per training sample size and per splitting iteration, such that the different models operated on the exact same data splits.

To carry out model selection, hyperparameters were handled in a data-dependent way by grid search <u>(Goodfellow et al. 2016)</u>, separately for each dataset, model, training set size, and splitting iteration. Hyperparamter grids were set up for each model based on best practices from the literature, and chosen empirically based on relative prediction accuracy on the validation set. Linear discriminant analysis has no hyperparameters to tune. For logistic regression and SVMs, the regularisation parameter C and gamma were distanced in powers of two. Regarding hyperparameter grids for kernel models, coefficients of polynomial and sigmoidal kernels were - 1, 0, or 1; and the degree of the polynomial kernel was set to two. After the best hyperparameter combination in the grid of candidate choices had been determined, the actual absolute accuracy was assessed on the unseen test set.

While our linear and kernel models completed hyperparameter selection and parameter estimation in the order of minutes, deep neural networks took orders of magnitude longer. For computational feasibility, deep neural networks were tuned only with regard to the L2 penalty on their parameters (0., 1e-3, 1e-5, 1e-7 for convolutional DNNs and 1e-4, 1e-3.75, …, 1e-0.5 for fully connected DNNs). Every architectural choice, such as whether or not to use dropout, filter or unit numbers per layer, learning rates, can be viewed as a form of hyperparameter of the model and could potentially be fine-tuned. A focus on tuning the regularisation strength is most often used as a middle ground between completeness and computational feasibility in the literature (Goodfellow et al. 2016). Please note that all candidate sets of hyperparameters choices were evaluated exclusively on validation set data. Additionally, all multi-class analyses were carried out as one-vs.-rest schemes for comparability (Hastie et al., 2001).

### Sample complexity analysis: Monitoring model performance with increasing sample size

To satisfy the core motivation behind the present investigation, we quantified how a given model’s prediction success scales as a function of the sample size at current disposal. For each model and dataset, we ran separate cross-validated prediction analyses for increasing steps of training set sizes: n = 100, 200, 500, 1,000, 2,000, 4,000, and 8,000. The resulting prediction estimates provided the basis to build a so-called “learning curve” (Goodfellow et al. 2016; Abraham et al. 2017). We will call the resulting relation between the available number of observations and prediction performance the empirical sample complexity of a given model on a particular dataset and a specific prediction goal.

Such learning curves of machine algorithms typically follow an inverse-power law. The accuracy often increases rapidly in the beginning and then slowly saturates (Cortes et al. 1994). These diagnostic assessments are typically further characterized by a saturation point and a saturation velocity. The saturation point provides a sense of the maximal performance that the prediction algorithm can achieve as the sample size keeps growing infinite to always more observations. The delineation of a model’s scaling of predictive performance is directly tied to the signal-to-noise ratio of the given dataset and the expressive capacity of the given model <u>(Goodfellow et al. 2016)</u>. The saturation velocity indicates how many observations are necessary to approach maximal prediction performance and is further tied to the complexity of the target function to be approximated by the model in the dataset. Different scaling behaviour of linear, kernel, and deep models can delineate the extent to which exploitable nonlinear structure is present in the data, and at what sample sizes such nonlinear information becomes practically relevant for realizing better predictions.

## Acknowledgments

We thank Parashkev Nachev, Robert Gray, Sami Hamdan, and Cembre Zor for insightful comments on a previous version of this manuscript. The authors declare no conflict of interest.

We thank the UK Biobank participants for their voluntary commitment and the UK Biobank team for their work in collecting, processing and disseminating these data for analysis. This research was conducted using the UK Biobank Resource under approved project 23827.

DB was funded by the Deutsche Forschungsgemeinschaft (DFG, BZ2/2-1, BZ2/3-1, and BZ2/4-1; International Research Training Group IRTG2150), Amazon AWS Research Grant (2016 and 2017), as well as the START-Program of the Faculty of Medicine (126/16) and Exploratory Research Space (OPSF449), RWTH Aachen. Simulations were performed with computing resources granted by RWTH Aachen University under project number “rwth0238”.

## Author Contributions

DB designed the study. MAS performed the experiments. DB, MAS, and BTTY analyzed the data. DB, JNK, KK, JMM, BR, MAS, JTV, and BTTY wrote the paper.

## Code and Data Availability

Source code is available at http://github.com/maschulz/deeperbrain. MNIST and Fashion data are freelyavailableat http://http://yann.lecun.com/exdb/mnist/ and http://https://github.com/zalandoresearch/fashion-mnist. UK BioBank brain-imaging data is available upon application at http://www.ukbiobank.ac.uk/register-apply/.

